# *ARID1A*-deficient cells promote endometrial carcinoma progression and dissemination by reprograming tumor microenvironment

**DOI:** 10.1101/2022.12.14.520528

**Authors:** Cristina Megino-Luque, Xavier Matias-Guiu, Núria Eritja

## Abstract

AT-rich interactive domain-containing protein 1A (ARID1A) loss-of-function mutation accompanied by a loss of ARID1A protein expression is frequently observed in endometrial carcinomas (EC). However, the mechanisms underlying this loss of ARID1A involved in the progression and dissemination of EC have been poorly studied and understood. Tumor microenvironment (TME) is reprogramed by cancer cells and have a high influence in aggressive behaviours of solid tumors. TME, and specially the stromal component, has a clear impact on the progression and aggressiveness of EC, but little is known about its activation, signalling and functions. Thus, the main aim of this study was to analyse the role of EC ARID1A loss in TME reprograming and its implication in endometrial tumor progression and aggressiveness. Here, using different endometrial *in vitro* and *in vivo* models, we show that endometrial tumor cells with altered expression of *ARID1A* promote the modulation and activation of the TME. We demonstrate that ARID1A depleted endometrial tumor cells induce aggressive behaviour in neighbouring endometrial tumor cells with wild-type ARID1A expression and the recruitment and activation of endometrial stromal cells (ESC). Interestingly, these pro-tumoral signals spread through the bloodstream supporting ARID1A wild-type EC cells aggressiveness. Finally, we show that ARID1A altered tumor cells display a different secretion profile and express significative high levels of chemokines in the TME. Taken together, our data demonstrate ARID1A depleted endometrial tumor cells reprograms TME promoting progression and dissemination of EC through chemokine secretion.

## Introduction

Endometrial cancer (EC) is the most frequent tumor of the female genital tract and the sixth most common cancer in women worldwide ^1^. Moreover, it is one of the few tumors whose incidence and mortality have increased in the recent years, probably related to the highest life span and obesity rates^2^. Classically, EC has been broadly classified into two groups based on clinical, pathological and molecular features: Type I or endometroid EC (EEC), composed by tumors with endometroid histology, representing approximately 85% of all EC and is usually associated with favorable prognosis; and type II or non-endometroid EC (NEEC) which account for 15% of all EC and is related to poor clinical outcomes^3,4^. In general, most EEC patients are diagnosticated in early tumor stages presenting a favorable prognosis and clinical courses. However, approximately 20% of EEC are diagnosed in high-grade stages, presenting more aggressive patterns, related to the appearance of recurrences and distal metastases, and being responsible for up to 39% of deaths from EEC^5,6^. Although this dualistic classification is broadly used, in 2013 a new molecular classification of EC stablished a new four EC genomic subtypes in bases of the genomic tumor profiles, detecting high frequencies of somatic mutations in several tumor suppressor genes such as ARID1A^7^.

ARID1A (At-rich interactive domain, also known as BAF250A) is the most important subunit of the SWI/SNF (SWitch/Sucrose Non Fermentable) chromatin remodeling complex compromising the DNA binding domain of this complex^8^. The SWI/SNF complex modulates gene transcription by remodeling nucleosomes in an ATP dependent manner, playing important roles in several cellular process such as tissue differentiation or proliferation^9^. The SWI/SNF complex is mutated in >20% of all human cancers, among which somatic alterations in ARID1A are the most frequent, leading to loss of function^10^. Specifically, ARID1A mutations occurs with high frequency in gynecological cancers, with a mutational frequency of 46,7% observed in low-grade EEC, rising to 60% in high-grade EEC^11^. Although the role of ARID1A in EC is not fully understand, recent findings suggest that in EEC, ARID1A participate in tumor progression rather than initiation^9,12,13^. Given the important role of ARID1A a better characterization of ARID1A lost effect in endometrial tumor progression is critical to improve ARID1A mutation EC treatments.

Tumors consist in a complex mixture of numerous cell types, soluble factors and extracellular matrix components (ECM), and their composition, interactions and polarization have a critical role in tumor progression and invasion. This complex tumor microenvironment (TME) is continually changing over the course of cancer progression, in response to evolving environmental conditions and oncogenic signals from growing tumors^14,15^. Cancer associated fibroblast (CAFs) represent one of the most abundant and plastic cell type in the TME and are involved in many tumor hallmarks such as tumor progression, dissemination and therapy responses through remodeling the ECM and signaling to the different cell types of the TME ^14–16^. The soluble factors released by cancer and stroma cells constitute a secretory profile that have a primordial role in tumor progression and malignant phenotypes acquisition^14,15^. In EC the TME have a pivotal role in malignant and metastatic endometrial tumor transition, and it is increasingly postulated that microenvironment-derived signals are necessary to drive EC progression^17,18^. However, the knowledge about TME influence in EC progression raising limited.

Herein, we report that ARID1A loss expression in EC cells is associated with more aggressive phenotypes in surrounding ARID1A-wildtype tumor cells and the activation of the endometrial stroma cells (ESC) promoting CAF phenotypes and leading to a reactive TME. This pro-tumoral TME supports endometrial tumor progression and dissemination, by dysregulation of the secretory tumor profile.

## Material and Methods

### Reagents and antibodies for western blot

The following reagents were used: anti-GAPDH (Abcam, 8245); anti-ARID1A (Cell Signaling,12354); anti-E-cadherin (BD Biosciences, 610181); anti-β-catenin (BD Biosciences, 610153); anti-N-cadherin (Santa Cruz, sc-7939); anti-Vimentin (BD bioscience, 550513); anti-SNAIL (Cell Signaling, 3879); anti-ZEB (Santa Cruz, sc-515797); anti-α-SMA (Santa Cruz, sc-53015); anti-FAP (Santa Cruz, sc-100528); anti-p-PDGFRa (Tyr754) (Santa Cruz, sc-12911); anti-p-NFKb-p50 (Ser337) (Santa Cruz, sc-101744); anti-p-NFKb-p65 (Ser536) (Cell signaling, 3033); anti-IL6 (Santa Cruz, sc-57315); anti-p-paxillin (Tyr118) (Cell signaling, 2541); anti-COL1A (Cell signaling 39952); anti-p-MLY2 (Ser19) (Cell signaling, 3671); Rhodamine-conjugated phalloidin (Sigma-Aldrich, P1951); anti-GFP (Sigma-Aldrich, # A-11122).

### Cell culture

Ishikawa 3-H-12 (IK) were purchased from Sigma-Aldrich (Sigma-Aldrich, 99040201), as well as MFE-296 cell line (Sigma-Aldrich, 98031101). Cells were grown in Dulbecco’s modified Eagle’s medium (DMEM; Sigma-Aldrich, 12007559) supplemented with 10% fetal bovine serum (FBS; Invitrogen, 10270106), 1 mmol/L HEPES (Sigma-Aldrich, H3375), 1 mmol/L sodium pyruvate (Sigma-Aldrich, P2256), 2 mmol/L L-glutamine (Sigma-Aldrich, C59202), 1% penicillin/streptomycin (Sigma-Aldrich, P4333) at 37C with saturating humidity and 5% CO2.

To generate Ishikawa 3-H-12 (IK) and MFE-296 sgRNA *ARID1A-*deficient cell lines, cells were infected whit the lentiviral plasmid encoding Cas9 and the sgRNA against *ARID1A*. Cells infected with viruses encoding the puromycin resistance gene were selected in 2 μg/ml puromycin.

### Isolation of Mouse endometrial epithelial cells and ESC

Isolation of mouse endometrial epithelial and stromal cells was performed as described previously with minor modification^19^. Briefly, after mice sacrifice uterus was dissected and washed with Hanks’ balanced salt solution and chopped in 3- to 4-mm-length fragments. Uterine fragments were digested with 1% trypsin (Invitrogen) in Hanks’ balanced salt solution (Invitrogen) for 1 hour at 4°C and 45 minutes at room temperature. Trypsin digestion was stopped by addition of Dulbecco’s modified Eagle’s medium (DMEM) containing 10% fetal bovine serum (Invitrogen). After trypsin digestion, epithelial sheets were squeezed out of the uterine pieces and separated from the stroma by applying gentle pressure with the edge of a razor blade. Epithelial sheets were washed twice with PBS and resuspended in 1 ml of DMEM/F12 (Invitrogen) supplemented with 1 mmol/L HEPES (Sigma-Aldrich), 1% penicillin/ streptomycin (Sigma-Aldrich), and Fungizone (Invitrogen) (basal medium). On the other hand, for endometrial stromal cells isolation, the stromal uterine pieces were cut with a scalpel blade into smaller fragments, washed with 1X PBS and centrifuged at 1000 rpm for 3 min. These fragments were then disrupted with a 1% collagenase IA solution (200mg/ml, Worthington Biochemical Corporation) in DMEM culture medium supplemented with FBS, and incubated for 2h at 37ºC with agitation at 700rpm. After this time the solution was filtered with a 40μM Steril Cell Strainer (Fisherbrand) and the isolated stromal cells were seeded in a p100 plate with stromal cell medium composed of DMEM/F12 supplemented with 4% FBS, 2 mmol/L sodium pyruvate, 1% penicillin/streptomycin and 0.5% Insulin-Transferrin-Sodium Selenite (ITS). After 24 h, the cells were washed with 1X PBS and the medium was changed to fresh medium. The cells were incubated at 37°C with saturated humidity and 5% CO2.

### Conditioned media (CM) collection

For the CM collection, the cells in question were seeded under their normal growth conditions and allowed to grow to 70% confluency. Once the confluence in question was reached the cultured cells were washed with 1X PBS and the desired fresh medium was added for harvesting. After 48h the CM were collected and centrifuged at 1000 rpm for 3 min. These CM were recovered and passed through a 22μM filter. Finally, the CM were stored at -80ºC until use.

### 3D spheroids co-cultures system of EC cell lines

Growth of human endometrial epithelial cell lines in cultures was performed as described previously with minor modification^19^. Briefly, cells were washed with Hanks Balanced Salt Solution HBSS (Thermo Fisher Scientific, 11520476) and incubated with trypsin-EDTA solution (Sigma-Aldrich, T4049) for 3 min at 37Cº. Trypsin activity was stopped by adding DMEM containing 10% FBS. Cells were centrifuged at 18 x g for 3 min. First, about 1,000 endometrial epithelial cells or endometrial stromal cells (those desired for each condition) were resuspended in 10 μl of Matrigel (BD Biosciences, 354234) per well of M96, generating a base that was allowed to polymerize at 37°C for 30 min. After this time, 1,500 endometrial GFP-labelled epithelial cells/well were resuspended in 100μl of DMEM/F12 medium (Sigma-Aldrich, 11580376) supplemented with 2% FBS, 1 mmol/L sodium pyruvate, 1% penicillin/streptomycin and 0.1% amphotericin B were seeded and incubated for 5 to 8 days at 37°C. Images were acquired using confocal microscopy (model FV1000; Olympus, Tokyo, Japan). Analysis was performed using ImageJ software and invasion index was calculated measuring the total area over which cancer cells had dispersed (including invading and non-invading cells) and the area of non-invading cells (1 – [non-invading area/total area]).

### Generation of organotypic cultures of EC cells and ESC

EC orthotopic culture system was set up as previously described ^20^ with some adaptations. Briefly, first of all 2.5×10^6^ mice ESC or CAFs were resuspended in 100μl of stromal cell medium (composed of DMEM/F12 supplemented with 4% FBS, 2 mmol/L sodium pyruvate, 1% penicillin/streptomycin and 0.5% ITS). This suspension was in turn dissolved in a mixture of 50% Collagen type I (4mg/ml; Thermo Fisher Scien-tific), 25% Matrigel, 6.25% FBS and stromal cell medium, generating a gel that was deposited in an M24 well and incubated for 1h at 37ºC with humidity saturation and 5% CO2. Next, 1 ml of stromal cell medium was carefully added to the well and incubated for 16h again at 37ºC with humidity saturation and 5% CO2. Then, the medium was aspirated very carefully and a suspension of 5.5 × 10^5^ tumor cells of interest resuspended in DMEM tumor cell medium was added to a final volume of 500μl. Again, incubation was repeated for 16h at 37ºC with humidity saturation and 5% CO2. Finally, the graft where the culture was mounted was prepared. For this, first 1ml of the mixture of 50% Collagen type I (4mg/ml; Millipore), 25% Matrigel, 6.25% FBS and stromal cell medium was prepared for each graft (Millipore) to be prepared. This mixture was deposited on the top of the graft and then incubated for 1h at 37ºC. After the hour of incubation, the gels were fixed with sterile PFA 4% for 2h at room temperature; washed three times with PBS 1X for 10min; and neutralized by adding tumor cell medium and incubating for 30min at 37ºC. Finally, these were carefully transferred to M6 wells. Once the grafts were prepared, the gels were placed on the grafts and tumor cell medium was added until reaching the base of the graft and 100μl of the mixture of 50% Collagen type I, 25% Matrigel, 6.25% FBS and stromal cell medium was added to each gel (on top of the tumor cell portion) for better preservation of the structure and orientation of the culture components. Finally, it was incubated for 6-7 days at 37ºC with humidity saturation and 5% CO2. After this period, for the analysis of the organotypic cultures, the cultures were fixed with formalin for 16h at 4ºC and HE was performed. Images were acquired using a Leica DMD170 microscopy. Analysis were performed using ImageJ and the invasion index was calculated by measuring the total area over which each EC cell had dispersed (including invading and non-invading cells) and the area of non-invading cells (1 – [non-invading area/total area]).

### Viral production, infection and *in vitro* transfection conditions

Oligonucleotides to produce plasmid-based sgRNA were cloned into the lentiviral lentiCRISPRv2 vector using BsmBI restriction sites. sgRNA targets sequence were: ARID1A-3 5’-CACCGTATGGCCAATATGCCACCTC. Target sequences were functional against both human *ARID1A* and mice *Arid1a* genes.

Production of viral particles was achieved by transfecting HEK-293 packaging cells with linear PEI (40 μM) in combination with lentiviral plasmids and helper plasmids (psPAX2 packaging and pMD2G envelope) at 1:1:1 ratio, respectively.

Four hours after transfection, packaging cells were cultured with DMEM supplemented with 10% FBS, 1 mmol/l HEPES (Sigma-Aldrich), 1 mmol/l sodium pyruvate (Sigma-Aldrich), 2 mmol/l L-glutamine (Sigma-Aldrich) and 1% of penicillin/streptomycin (Sigma-Aldrich) for 3-4 days; afterwards the medium containing the viral particles was collected, centrifuged for 10 min at 200 x g and filtered thought a 0.45 μM filter (Millipore, Temecula, CA, USA) and concentrated using Vivaspin concentrators (Sartorius Stedim Biotech GmbH, Gottingen, Germany). The concentrated medium containing lentiviral particles was added to the medium of the pre-plated cells. Cells were incubated for 24 h. After this period, the medium was replaced with fresh medium and cells were grown regularly to allow phenotypic expression.

### Genetically modified mouse models

The in vivo studies complied with Law 5/1995 and Act 214/1997 of the Regional Government (Generalitat de Catalunya) and EU Directive EEC 63/2010 and were approved by the Ethics Committee on Animal Experiments of the University of Lleida and the Ethics Commission in Animal Experimentation of the Generalitat de Catalunya. *Cre-ER*^*T*^ (B6. Cg-Tg(CAG-Cre/Esr1*5Amc/J) and *Pten* ^f/f^ (C;129S4-Ptentm1Hwu/J) mice were obtained from the Jackson Laboratory (Bar Harbor, ME, USA). *Arid1a* ^f/f^ mice were a kind gift from Dr. I. Lei. Mice bearing floxed *Arid1a* allele in which exon 9 of *Arid1a* gene is flanked by loxP sites have been described before^21^. Cre:ERT^+/−^PTEN^f/f^ mice were generated as described previously^22^ .*Arid1a*^*f/f*^ mice were backcrossed for five generations with C57BL/6 mice before being crossed with *Cre-ER*^*T*^PTEN^f/f^ transgenic strains to generate epithelial cell-specific deletion of Arid1a. Mice were genotyped by earmarking and DNA was isolated from tail tissue in proteinase K lysis buffer. PCR was carried out with GoTap polymerase (Promega, Madison, WI, USA) using different pairs of primers for each gene. *Cre-ER*^*+/−*^ forward primer 5′-ACGAACCTGGTCGAAATCGTGCG-3′ and reverse primer 5′-CGGTCGATGCAACGAGTGATGAG-3′; *Pten*^*f/f*^ forward primer 5′-CAAGCACTCTGCGAACTGAG-3′ and reverse primer 5′-AAGTTTTTGAAGGCAAGATGC; *Arid1a*^*f/f*^ forward primer 5′-GGCTCTGCCATAAAGCGATCC-3′ and reverse primer 5′-CTCACAAATCTAACCGAGGCCAC-3′.

### Tamoxifen administration Tamoxifen

As previously described^22^, tamoxifen (Sigma-Aldrich T5648, St Louis, MO) was dissolved in 100% ethanol at 100 mg/ml. Tamoxifen solution was emulsified in corn oil (Sigma-Aldrich C8267) at 10 mg/ml by vortexing. To induce PTEN and ARID1A deletion, adult mice (4-5 weeks old) were given a single intraperitoneal injection of 0.5 mg of tamoxifen emulsion (30-35 μg per mg body weight).

### Immunohistochemical study

After sacrifice, mice uteri were excised, flushed with PBS, fixed in 10% neutral-buffered formalin and embedded in paraffin. Mice uterus blocks were sectioned and thickness of 3μm, dried for 1 hour at 65ºC before pre-treatment procedure of deparaffinization, rehydration and epitope retrieval in the Pre-Treatment Module, PT-LINK (Agilent Technologies-DAKO) at 95 °C for 20 min in 50 × Tris/EDTA buffer, pH 9. Before staining the sections, endogenous peroxidase was blocked. The antibodies used were anti-ARID1A (1:500 dilution, Abcam, ab182561) and Ki-67 (1:100 dilution, Abcam, ab16667,). After incubation, the reaction was visualized with the EnVisionTM FLEX+ rabbit (Linker) Detection Kit (Agilent Technologies-DAKO) for ARID1A and the secondary antibody polyclonal goat anti rabbit IgG/Biotin (1:200 dilution, Jackson Immunoresearch, 111-065-144) plus Streptavidin/HRP (1:400 dilution, Agilent Technologies-DAKO, P0397)) for Ki67, using diaminobenzidine chromogen as a substrate. Sections were counterstained with hematoxylin. Appropriate negative controls including no primary antibody were also tested.

Immunohistochemical results were evaluated by following uniform pre-established criteria. Immunostaining was graded semi-quantitatively by considering the percentage and intensity of the staining. A histological score was obtained from each sample and values ranged from 0 (no immunoreaction) to 300 (maximum immunoreactivity). The score was obtained by applying the following formula, Histoscore=1 × (% light staining) +2 × (% moderate staining) +3 × (% strong staining). To support the scoring of immunohistochemistry, a digital slide scanner [Nuclear Quant Module, Pannoramic 250 FLASH II 2.0 (3D HISTEC) was used and the percentage of positive cells was determined.

### Transwell assay

1×10^4^ cells per well were plated in the upper chamber of the Transwell (8μm pore, Falcon) previously coated with Matrigel in serum-free medium at a density. As a chemoattractant were used collected CM or 10^6^ cells of interest previously planted at the well base during 16h. After 48 h, cells were fixed with paraformaldehyde 4% and stained with Hoechst (5μg/ml). Finally, cells were pictured with an epifluorescence microscope (Leica), before and after a cotton swap. Results were analyzed to obtain the percentage of invasive cells using the software Image J.

### Wound healing assay

To assess cell migration by wound healing assays, 6×10^4^ cells of interest were seeded per well in a 24-well plate and triplicates of each condition were performed. When the cells had reached 70% confluence, a wound was made with the end of a 200μl tip and a wash was performed with 1X PBS to remove any remaining cells floating in the medium and fresh medium or the medium of interest to be analyzed was added. For each well, three photographs at different heights were taken at time 0, which were replicated after 48h of incubation at 37°C with humidity saturation and 5% CO2. Finally, the analysis of the % of closed wound was calculated using Image J software.

### Contractibility assay

To assess contractibility, 250,000 stromal cells of interest per gel were resuspended in 100μl of a mixture of 2mg/ml collagen and Matrigel in a 1:1 ratio. These gels were placed in wells of M24 plates and incubated for 1h at 37°C with humidity saturation and 5% CO2. After this time 500μl of stromal cell medium was added and allowed to incubate for 5-6 days. Shrinkage of the collagen gels was monitored by scanning the plates. For analysis of the results, the relative well and gel area was measured using Image J software. Percentage of contractibility was calculated employing the formula 100 × (well area − gel area) / well area.

### Total RNA extraction, reverse transcriptase-PCR and quantitative real-time

For RT-qPCR, total RNA was extracted from the 3D or 2D cultures using the RNeasy Total RNA kit (Qiagen, Valencia, CA, USA). The cDNA synthesis and RT-qPCR steps were performed simultaneously using the pPCRBIO Probe 1-Step Go kit (PCR Biosystems). The program used for the process starts with heating the samples at 45°C for 10min, then the samples are raised to 95°C for 2min and finally 40 cycles are repeated at 95°C for 5sec and 60°C for 20sec, using a CFX96TM thermal cycler (BioRad). Relative mRNA expression levels were calculated using the 2^ΔΔCt^ method and are presented as ratios to the housekeeping gene *GAPDH*. Taqman technology from Applied Biosystems was used for real-time RT-qPCR analyses. Probes: mouse *Vegf* (Mm00437304_m1); mouse *Il6* (Mm00446190_m1); mouse *Tgfb1* (Mm00441724_m1);; mouse *Gapdh* (Mm99999915_g1).

### Immunofluorescence assay

2D or 3D cultures were fixed with paraformaldehyde 4% for 15 min at room temperature and washed twice with PBS. Depending on primary antibody, cells were permeabilized with 0.2% Triton X-100 in PBS for 10 min or with 100% methanol for 5 min. Next, cultures were incubated overnight at 4ºC with the indicated diluted primary antibodies: anti-α-SMA (1:200, Triton X-100; Santa Cruz, sc-53015); anti-FAP (1:200, Triton X-100; Santa Cruz, sc-100528); anti-COL1A (1:200, Triton X-100; Cell signaling 39952); anti-p-MLY2 (Ser19) (1:200, Triton X-100; Cell signaling, 3671); Rhodamine-conjugated phalloidin (1:500, Triton X-100; anti-CXCL16 (1:200, Triton X-100; Santa Cruz, sc-514363); anti-p-paxillin (Tyr118) (Cell signaling, 2541); Sigma-Aldrich, P1951); anti-GFP (1:1000, Triton X-100; Sigma-Aldrich, # A-11122). Then, cells were washed twice with PBS and incubated with PBS containing 5μg/mL of Hoechst 33342 and a 1:500 dilution of secondary anti-mouse Alexa Fluor 546 (Invitrogen, A11005) and Alexa Fluor 488 (Invitrogen, A11029) or anti-rabbit antibodies Alexa Fluor 594 (Invitrogen, R37119) and sAlexa Fluor 488 (Invitrogen, A11034) for 4 h at room temperature. Immunofluorescence stains was visualized and analyzed using confocal microscopy (model FV1000; Olympus, Tokyo, Japan) with the 10x, 20x and the oil-immersion 60x magnification objectives. Analysis of obtained images was performed with Fluoview FV100 software (Olympus, Tokyo, Japan) and ImageJ.

### Western blotting

Western blotting analysis were performed as described previously^23^. Briefly, cells were washed with cold PBS and lysed with lysis buffer (Tris HCl 50mM, NaCl 150mM, 1% de Tritón X-100, 0,1% de SDS, EDTA 1mM) and sonicated. Protein was quantified by Lowry (BioRad) and equal amounts of proteins were subjected to SDS-polyacrylamide gel electrophoresis and transferred to polyvinylidene difluoride membranes. Then membranes were blocked by incubation with TBST (20 mM Tris-HCl (pH 7.4), 150 mM NaCl and 0.1% Tween 20) plus 5% of non-fat milk. Membranes were incubated with the primary antibodies overnight at 4 °C and for 1h at room temperature with secondary horseradish peroxidase (1:10 000 in TBST). Signal was detected with SuperSignal West Femto Trial Kit (Thermo Scientific). Band analysis and densities were determined by using Image Lab 4.0.1 software (Bio-Rad laboratories, Richmond, CA, USA).

### Chemokine array

For the study of the chemokine secretion profile, the Human Chemokine Array G1 (bioNova, RayBio®; AAH-CHE-G1-4) was used. The first step was to place the chip containing the chemokine probes to be detected under a laminar flow hood for 2h to achieve complete drying. Subsequently, 100μl of blocking buffer was added and incubated for 30min at room temperature. After this time the buffer was aspirated, 100μl of the indicated samples were added to each sub-array and left to incubate for 16h at 4ºC under agitation. Then, a process of consecutive washes with different wash buffers (provided by the kit) was carried out to ensure the elimination of possible non-specific signals. Finally, for detection, 70μl per well of a mixture of biotin-conjugated primary antibodies was added and incubated for 16h at 4ºC with constant agitation. Subsequently, after a sequence of washes, 70μl per well of a fluorophore-labeled streptavidin solution was added (so it was always protected from light) and incubated for 2h at room temperature. Finally, several consecutive washes with different washing buffers were performed and finally it was left to dry under a laminar flow cabinet. Finally, analysis was performed using a laser scanner (Innopsys ‘InnoScan®) using the cy3 or “green” channel with an excitation frequency of 532 nm. The fluorescence intensity is proportional to the amount of cytokine in the test sample. Finally, for the heatmap and principal component analysis (PCA), the data were analyzed and represented using ClustVis123 software. For the clustering of the heatmap, the Euclidean distance was used for hierarchical grouping and the data grouping method was based on the average. For the analysis of the relative amount of each cytokine in the different samples, the data were analyzed and represented using GraphPad Prism v8 software.

### Statical analysis

All experiments were carried out at least 3 independent replicates, with at least 3 technical replicates. Statistical analysis of the data was carried out with *GraphPad Prism v8* software to determine if there were significant differences between the results of the different samples. The results were presented as the mean ± standard deviation (SD). All statistical analyses were set at a statistical significance level of p=0.05, correcting the p-values by the multiple comparisons test when appropriate. The data were analyzed using the most appropriate statistical test for each time point and the p-values are indicated in the figures.

In summary, for the analysis of means, statistical significance was verified by applying the Kolmogorov-Smirnov normality test, followed, in the case of parametric data, by Student’s t-test or the One-way ANOVA test (depending on whether the comparison was between only two variables or between more than two, respectively) and multiple comparisons by Tukey’s test. In the case of nonparametric data, the Mann-Whitney test was used for the comparison of two variables or the Kruskal-Wallis test for more than two. For the evaluation of the relationship of a main variable on the rest of the independent variables as well as on the rest of the main variables, the Two-way ANOVA test followed by a Bonferroni post-hoc test was performed. For all column analyses, contingency table comparisons, Fisher’s F test was used to compare variances. P values of less than 0.05 were considered statistically significant. Where the following symbols were used to describe statistical significance: *P < 0.05; **P < 0.01; ***P < 0.001; n.s., non-significant.

## Results

### ARID1A perturbed EC cells promotes more aggressive phenotypes in surrounding ARID1A wild-type EC cells

In previous studies we have reported that ARID1A alterations in stablished EC cells promote EMT phenotypes, inducing higher migratory and invasive tumor capacities, conferring more aggressive tumor phenotypes^23^. Tumors consist in a complex microenvironment composed of infiltrating and resident host cells, secreted factors and extracellular matrix^14,15,24^. Given these soluble mediators have and important role in tumor aggressiveness promoting migration and invasion^25^, we hypothesized that ARID1A downregulated cells could be inducing signals that drive to ARID1A wild-type cells alterations.

To test this hypothesis, we derived conditional medias (CM) from IK and MFE-296 ECC cell lines infected with lentiviruses carrying sgRNA against *ARID1A* (lentiCRISPRv2-sgRNA ARID1A-3) (CM-3) or with the empty vector (lentiCRISPRv2) (CM-V). CM-3 significantly promoted the migration of ARID1A wild-type cells, compared to CM-V treatments, as determined by wound healing assay (Fig. 1A and Supp. Fig. 1A). In support of these results, we observed that in transwell assays and three dimensional (3D) co-cultures models, ARID1A downregulated EC cells promoted invasion of ARID1A wild-type tumor cells (Fig. 1B-C and Supp. Fig. 1B). Moreover, when we analyzed the status of the EMT phenotypes in these cells, by immunoblotting assays, we observed that CM-3 elicited a down-expression on of several epithelial markers (such as E-cadherin or β-catenin) and an up-regulation of several mesenchymal markers (such as N-cadherin or Vimentin) and pro-EMT transcription factors (like SNAIL1 or ZEB) in ARID1A wildtype EC cells (Fig. 1D and Supp. Fig. 1C). Consistently with these results, co-culture assays showed that ARID1A downregulated IK cells significantly induced the expression of the mesenchymal marker Vimentin in ARID1A wild-type cells labelled with GFP (Fig. 1E). Further investigating the signal induction of ARID1A downregulated EC cells in ARID1A wild-type cells, we performed wound healing migration and transwell invasion assays in parenteral cells after treatments of 72h with the corresponding conditional medias. Consistent with the previous results, cells had significant increase in migratory and invasive abilities when they were previously treated with CM-3 comparing with CM-V treatments (Fig. 1F-G and Supp. Fig. 1D-E), demonstrating sustained activation in ARID1A wild-type cells by ARID1A altered cells.

**Figure 1.**
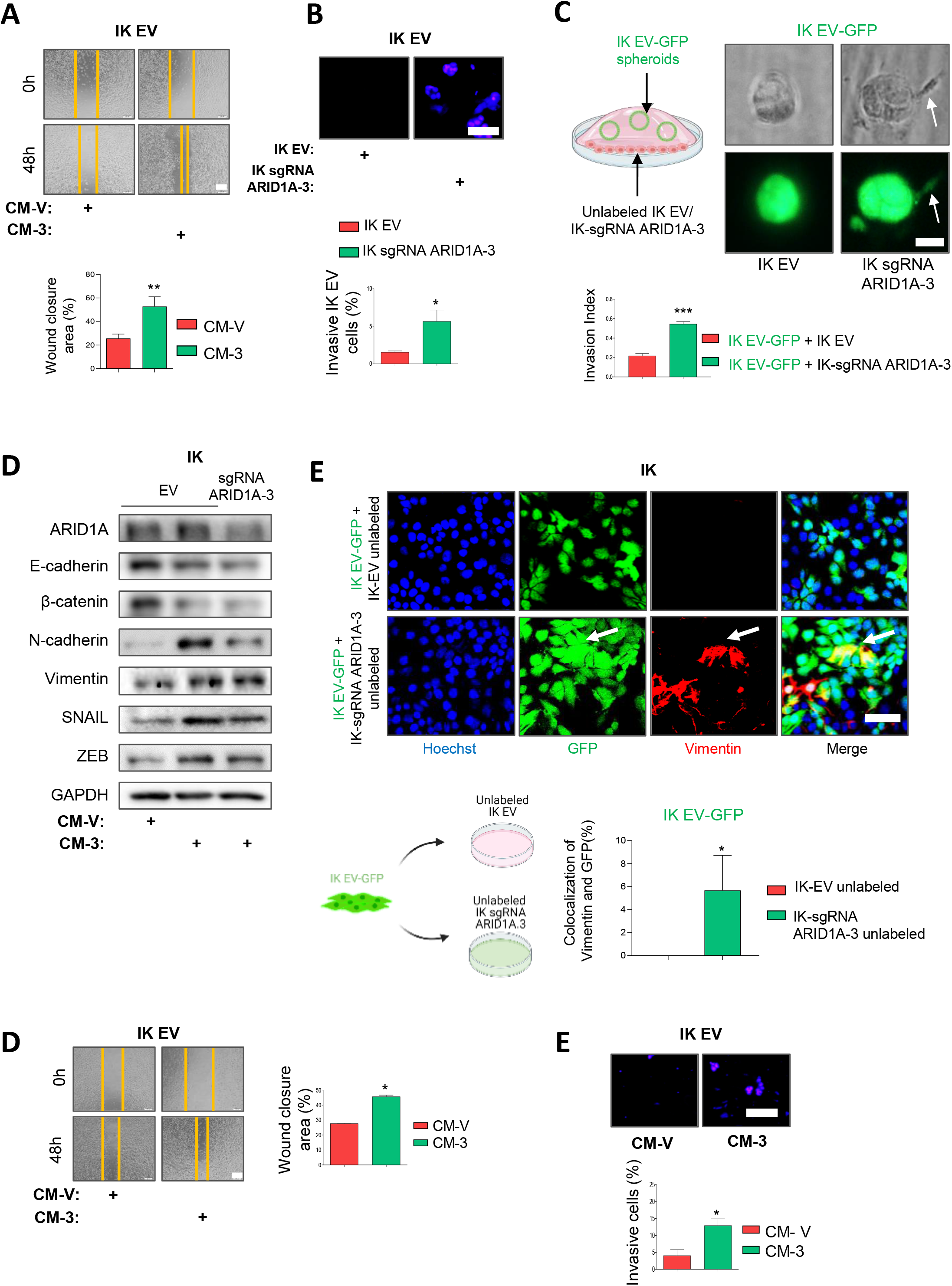
ARID1A perturbed EC cells promotes more aggressive phenotypes in surrounding ARID1A wild-type EC cells. **A)** Representative images at time 0 and 48 h after scratch, of wound-healing assay performed in IK cells infected with the EV treated with conditioned media collected from IK cells infected with lentiviruses carrying sgRNA against *ARID1A* (lentiCRISPRv2-ARID1A-3) (CM-3) or EV (CM-V). Bottom blot shows quantification of wound closure area between the indicated time. Scale bars: 200 μm. **B)** Representative images of nuclear Hoechst staining of transwell invasion assay after the cotton swab in IK control cells chemoattract by IK cells infected with lentiviruses carrying sgRNA against *ARID1A* (lentiCRISPRv2-ARID1A-3) or EV for 48 hours (upper panel), and quantification of Matrigel® invasive cells (bottom plot). Scale bars: 50 μm. **C)** Representative diagram (left) and panel of phase contrast images and immunofluorescence against GFP of 3D of IK cells infected with GFP and EV (IK EV-GFP) co-cultured with IK cells without labelled with GFP and infected or not with sgRNA against *ARID1A* (lentiCRISPRv2-ARID1A-3) (Right). Bottom plot shows invasive index quantification of IK EV-GFP glands in Matrigel. Scale bars: 25 μm. **D)** Western blot analysis of ARID1A, E-cadherina, β-catenina, N-cadherina, vimentina, SNAIL and ZEB in IK cells infected with the EV treated with CM-3 or CM-V for 48 hours. GAPDH was used as a loading control. **E)** Representative images at time 0 and 48 h after scratch, of wound-healing assay performed in IK cells infected with the EV previously treated for 48h with conditioned media collected from IK cells infected with lentiviruses carrying sgRNA against *ARID1A* (lentiCRISPRv2-ARID1A-3) (CM-3) or EV (CM-V). Right blots show quantification of wound closure area between the indicated time. Scale bars: 200 μm. **F)** Representative images of nuclear Hoechst staining of transwell invasion assay after the cotton swab in IK EV cells pretreated for 48 with conditioned media collected from IK cells infected with lentiviruses carrying sgRNA against *ARID1A* (lentiCRISPRv2-ARID1A-3) (CM-3) or EV (CM-V). (Left panel), and quantification of Matrigel® invasive cells (right plot). Scale bars: 50 μm. Graph values are the mean and error bars represented as mean ± S.E.M. Statistical analysis was performed using unpaired 2-tailed *Student t-test* analysis. * *p* < 0.05; ** *p* < 0.01; \*\*\**p* < 0.001. EV, empty vector.

### Pro-tumoral signals from ARID1A perturbed EC cells are spread through plasma, *in vivo*

Soluble factors secreted by the primary tumor, in addition to playing an important role in early stages of the metastatic cascade, also play an important role in more advanced stages of the metastatic cascade, fostering processes such as intravasation, circulation, extravasation and the generation of premetastatic niches^26^. This encouraged us to hypostatize that the observed pro-tumoral signals from ARID1A perturbed EC cells are extrapolated to plasma in order to promote tumor aggressiveness. To test this, *Cre:ERT; Pten f/f; Arid1a +/+* or *Cre:ERT; Pten f/f; Arid1a f/f* mice were injected intraperitoneally with one single dose of tamoxifen. After 6 weeks, all mice had developed endometrial tumors and their plasmas were collected (Fig. 2A). ARID1A loss expression in the uterus was assessed by immunohistochemistry (Fig. 2B). Next, we treated IK or MFE 296 ARID1A wild-type cells with the collected plasmas, observing than plasmas collected from *Cre:ERT; Pten f/f; Arid1a f/f* mice (P-f) induced higher migratory e invasive capacities than plasmas collected from *Cre:ERT; Pten f/f; Arid1a +/+* mice (P-+) in this ARID1A wild-type EC cells (Fig.2 C-D). In support of this, we observed that treatments with P-f, also promoted the acquisition of EMT phenotypes in IK or MFE 296 ARID1A wild-type cells, enhanced the protein expression of mesenchymal markers such as N-cadherin, vimentin, SNAIL or ZEB and decreased the protein expression of epithelial markers such as E-cadherin or β-catenin (Fig. 2 E). Taken together, these finding highlight that ARID1A perturbed EC cells induce pro-tumoral signals in mice plasmas.

**Figure 2.**
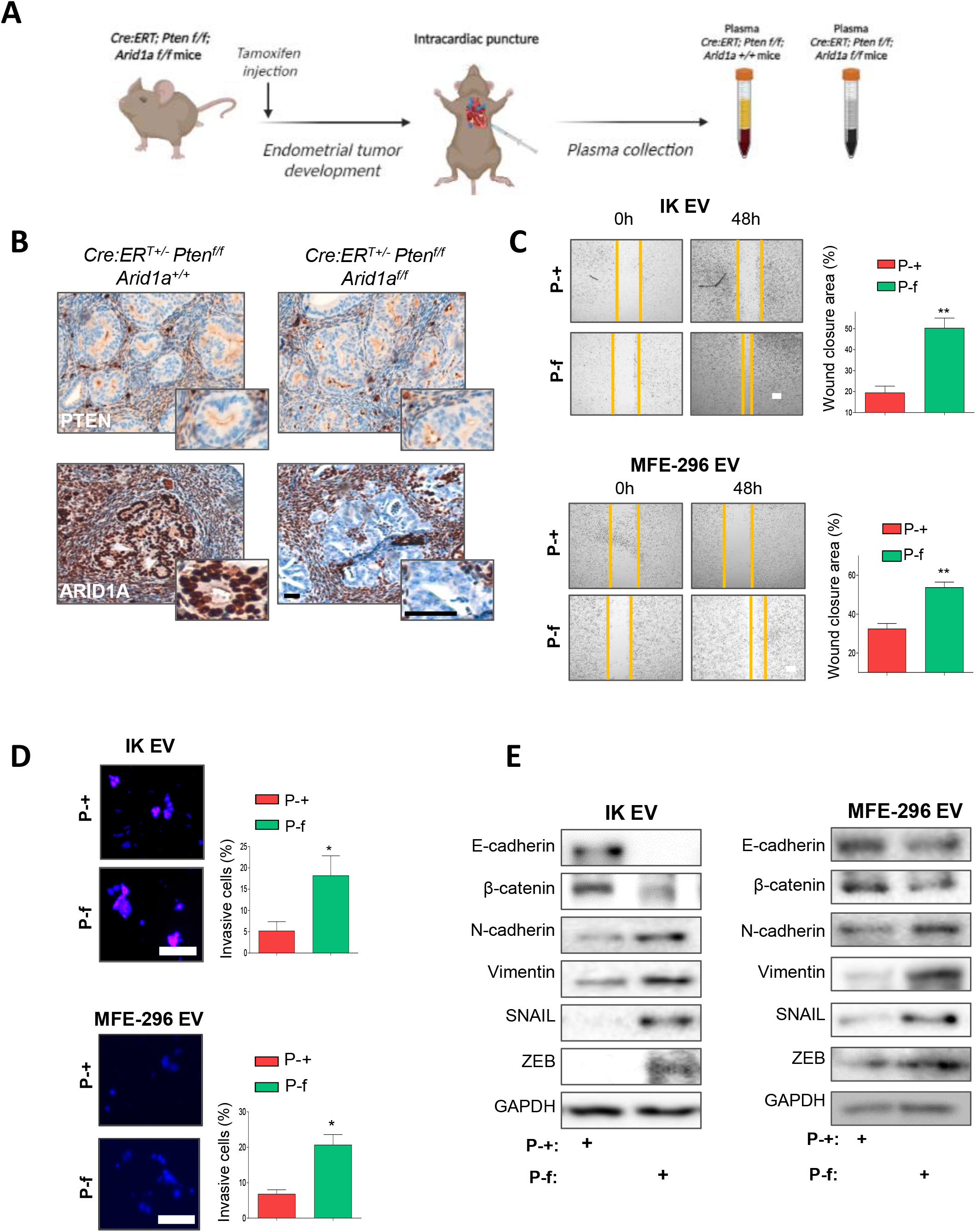
Pro-tumoral signals from ARID1A perturbed EC cells are spread through plasma, *in vivo*. **A)** Schematic representation of the experimental design for plasma collection. Briefly, *Cre:ERT; Pten f/f; Arid1a +/+* or *Cre:ERT; Pten f/f; Arid1a f/f* mice were weaned 3 weeks after birth and, after 5 weeks from weaning, were injected with a single dose of Tamoxifen to achieve double ablation of *Pten* and *Arid1a* in endometrial cells. After 6 weeks, all mice had developed endometrial tumors and plasma from *Cre:ERT; Pten f/f; Arid1a +/+* (P-+) or *Cre:ERT; Pten f/f; Arid1a f/f* (P-f) mice were collected. **B)** Representative images of ARID1A and PTEN immunohistochemistry in consecutive sections of uterus from *Cre:ERT; Pten f/f; Arid1a +/+* or *Cre:ERT; Pten f/f; Arid1a f/f* mice with EC. **C)** Representative images at time 0 and 48 h after scratch, of wound-healing assay performed in IK or MFE-296 cells infected with the EV treated with P-+ or P-f. Right blot shows quantification of wound closure area between the indicated time. Scale bars: 200 μm. **D)** Representative images of nuclear Hoechst staining of transwell invasion assay after the cotton swab in IK or MFE-296 EV cells treated for 48 with P-+ or P-f (left panel), and quantification of Matrigel® invasive cells (right plot). Scale bars: 50 μm. **E)** Western blot analysis of ARID1A, E-cadherina, β-catenina, N-cadherina, vimentina, SNAIL and ZEB in IK or MFE-296 cells infected with the EV treated for 48 with P-+ or P-f. GAPDH was used as a loading control. Graph values are the mean and error bars represented as mean ± S.E.M. Statistical analysis was performed using unpaired 2-tailed *Student t-test* analysis. * *p* < 0.05; ** *p* < 0.01. EV, empty vector.

### ARID1A depleted EC cells promotes endometrial stromal cells (ESC) activation

Cancer associated fibroblasts (CAF) are the major stromal cell type and one of the most active components within the complex and evolving tumor ecosystem^15,27,28^. Given it have been shown that tumor cell signals regulates stromal cell activation and functions^29,30^, we hypothesized that signals from ARID1A depleted EC cells promote ESC activation. To test this hypothesis, ESC isolated from wild-type mice were stimulated with CM treatments collected from IK EC cells (CM-V or CM-3). Notably, the ESC pre-treated with CM-3 showed higher elongated morphologies typical of CAF, compared to CM-V treated ESC (Fig. Supp 2A). In addition, CM-3 treatments promoted in ESC the characteristic ability of CAFs to contract collagen gels, increasing their matrix remodeling capabilities^31^ (Fig. 3A). In agreement with these results, organotypic co-culture model shown in figure 3B revealed that matrices generated by ESC previously treated with CM-3 were rendered significantly more permissive for the invasion of EC cells than the control, demonstrating also an increased pro-tumoral function in these activated ESC. Analysis by qRT-PCR, immunoblotting or immunofluorescence assays of several markers associated to stromal activation and CAF signatures (such as α-SMA, FAP, IL6, VEGFA, TFGβ1, COL1A and phosphorylated PDGFRα, NFKβ50, NFKβ65, paxillin and MLY2)^31^, corroborated that CM-3 treatments promoted ESC activation more than CM-V treatments (Fig. 3C-E). Moreover, immunofluorescence assay against F-actin, p-paxillin and p-MLY2 revelled that CM-3 induced cytoskeletal changes typical of CAF phenotypes, with more pronounced F-actin stress fibres, increased levels of p-paxillin forming focal adhesions and a strong association of p-MYL2 with actin stress fibres (Fig. 3E). Similar results were obtained in ESC treated with the plasmas collected from *Cre:ERT; Pten f/f; Arid1a +/+* or *Cre:ERT; Pten f/f; Arid1a f/f* mice with endometrial tumours, where P-f treatments induced collagen matrix contractibility (Fig. Supp. 2B) and higher expression levels of markers of CAF phenotypes such as α-SMA, FAP, IL6, VEGFA, TFGβ1, COL1A and phosphorylated PDGFRα, NFKβ50 and NFKβ65 (Fig. Supp. 3C-E).). Collectively, these findings emphasise the relationship between ARID1A depletion in endometrial tumour cells and the establishment of a reactive tumor stroma.

**Figure 3.**
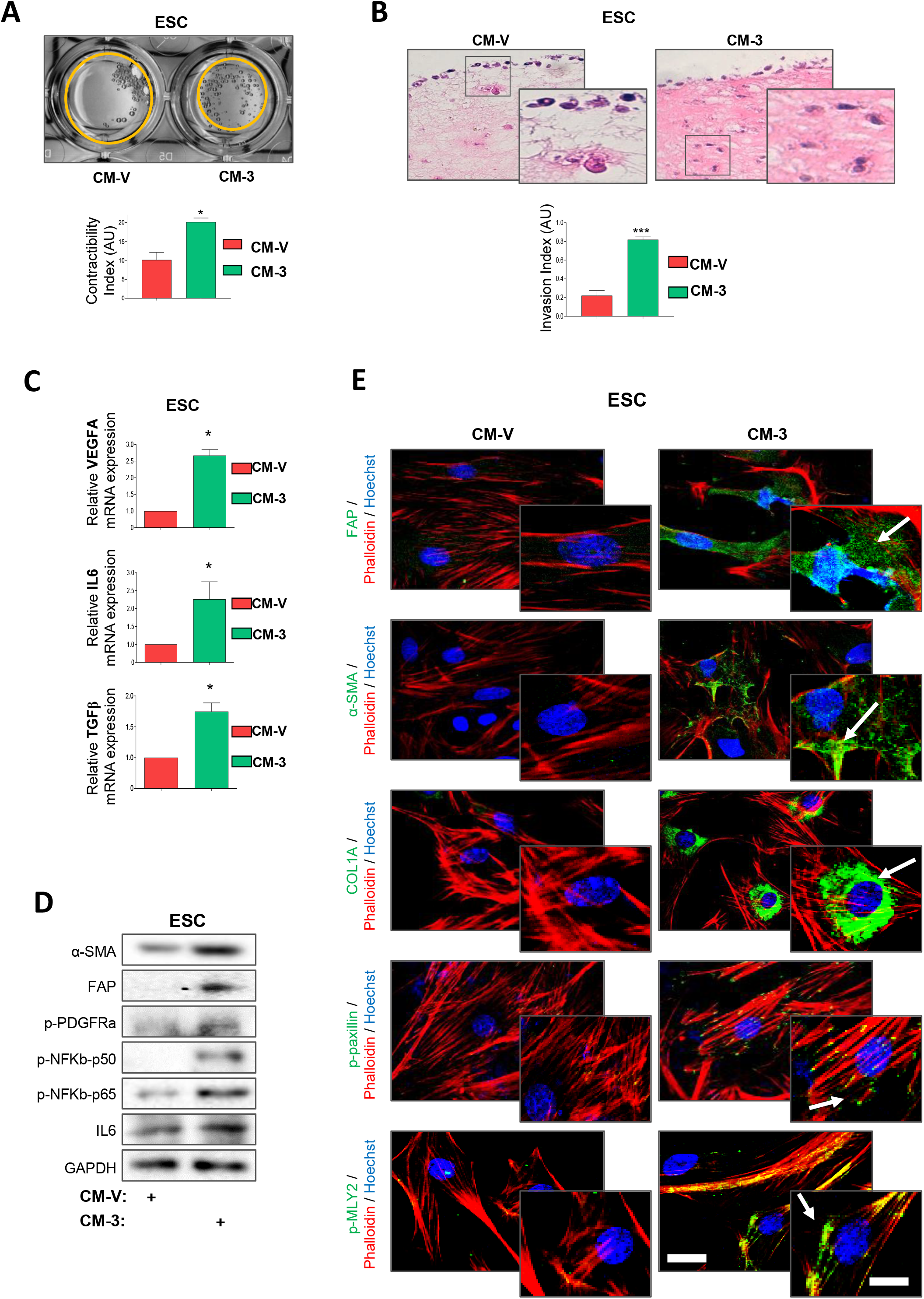
ARID1A depleted EC cells promotes endometrial stromal cells (ESC) activation and modulates their pro-tumorigenic functions. **A)** Upper, representative macroscopic images of collagen gels remodelled by ESC previously treated for 48h with CM collected from IK cells infected with lentivirus carrying sgRNA against ARID1A (lentiCRISPRv2 ARID1A-3) (CM-3) or empty vector (CM-V). Bottom, histogram representing the quantification of the contractility index. **B)** Representative Haematoxylin-Eosin (H&E) staining images of organotypic cultures showing invasion of IK EV cells into collagen matrices remodelled by ESC previously activated for 48h with CM-3 or CM-V (left), and blot showing quantification of IK EV cells invasion Index (right). Scale bars: 100 μm. **C)** RT-qPCR of *Vegfa, Il6* and *Tgfβ1* mRNA expression of ESC treated with CM-3 or CM-V for 48h. **D)** Representative immunoblotting images of α-SMA, FAP, p-PDGFRα (Tyr754), p-NFκβ-p50 (Ser337), p-NFκβ-p65 (Ser536) and IL6 protein expression in ESC treated with CM-3 or CM-V for 48h. GAPDH was used as a loading control. **E)** Representative immunofluorescence images of the stromal activation markers FAP, α-SMA, COL1A, p-paxillin (Tyr118) and p-MYL2 (Ser19) in ESC treated with CM-3 or CM-V for 48h. Phalloidin and Hoechst were used as actin cytoskeleton and nucleus markers respectively. GAPDH was used as a loading control. Graph values are the mean and error bars represented as mean ± S.E.M. Statistical analysis was performed using unpaired 2-tailed *Student t-test* analysis. * *p* < 0.05; ** *p* < 0.01; \*\*\**p* < 0.001. EV, empty vector.

**Figure 4.**
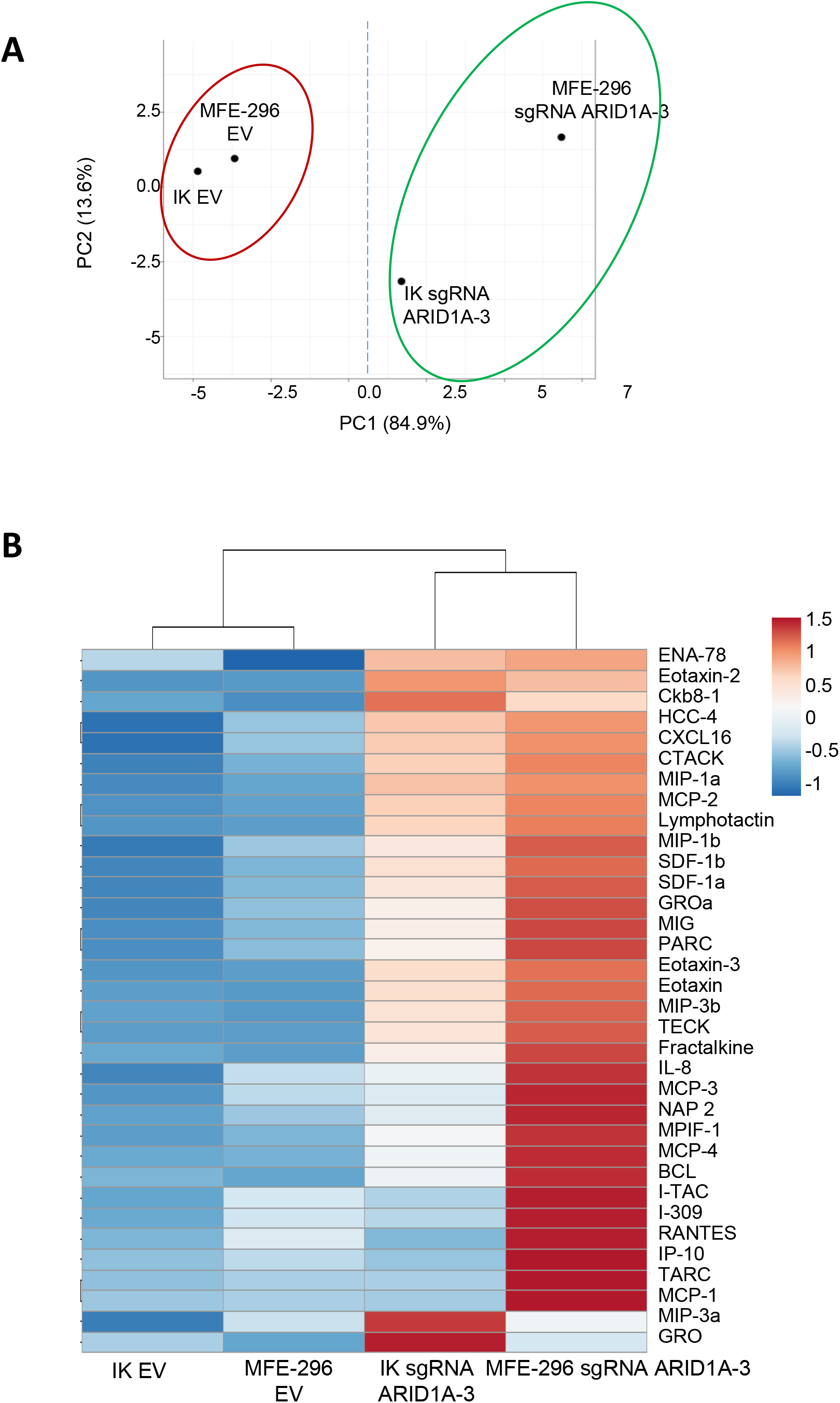
ARID1A depleted EC cells induced increased chemokine levels secretion. **A)** Principal component analysis (PCA) of chemokine array results showing change in the secreted profile of CM collected after 48h of IK or MFE-296 cells infected with lentivirus carrying sgRNA against ARID1A (lentiCRISPRv2 ARID1A-3) (CM-3) or empty vector (CM-V). **B)** Heatmap showing in hierarchical clusters the normalized levels of the chemokine array analysis performed with CM-3 and CM-V from IK and MFE-296 cells.

### ARID1A depleted EC cells induced increased chemokine levels secretion

Soluble mediators from the complex tumor microenvironment can foster tumor aggressiveness^24^. Given the obtained results, we hypothesized that these are, at least initially, driving by paracrine signaling through the expression of soluble factors due to ARID1A loss. As chemokines have a dramatic role facilitating cross-talk between tumor cells and their TME, we employed a chemokine array to study potential differences in the secretome profiles of EC cells with ARID1A wild-type or depleted expression. We analyzed the abundance of 38 soluble chemokines in CM-V and CM-3 from IK and MFE-296 EC cells. Principle component analysis (PCA) and unsupervised hierarchical clustering of chemokine expression levels, highlighted that EC cells engaged discrete chemokine secretory programs depending on ARID1A downregulation with the sgRNA ARID1A-3 infection (Fig. 5A-B). Specifically, we observed that conditional media from IK and MFE-296 ARID1A downregulated cells displayed moderately elevated levels of all analyzed tumor chemokines compared to parenteral cells with ARID1A wild-type expression (Fig. 5B). Taken together these results support the idea that ARID1A depleted tumor cells promotes more aggressive tumor phenotypes and stromal activation by chemokine secretion.

**Figure 5.**
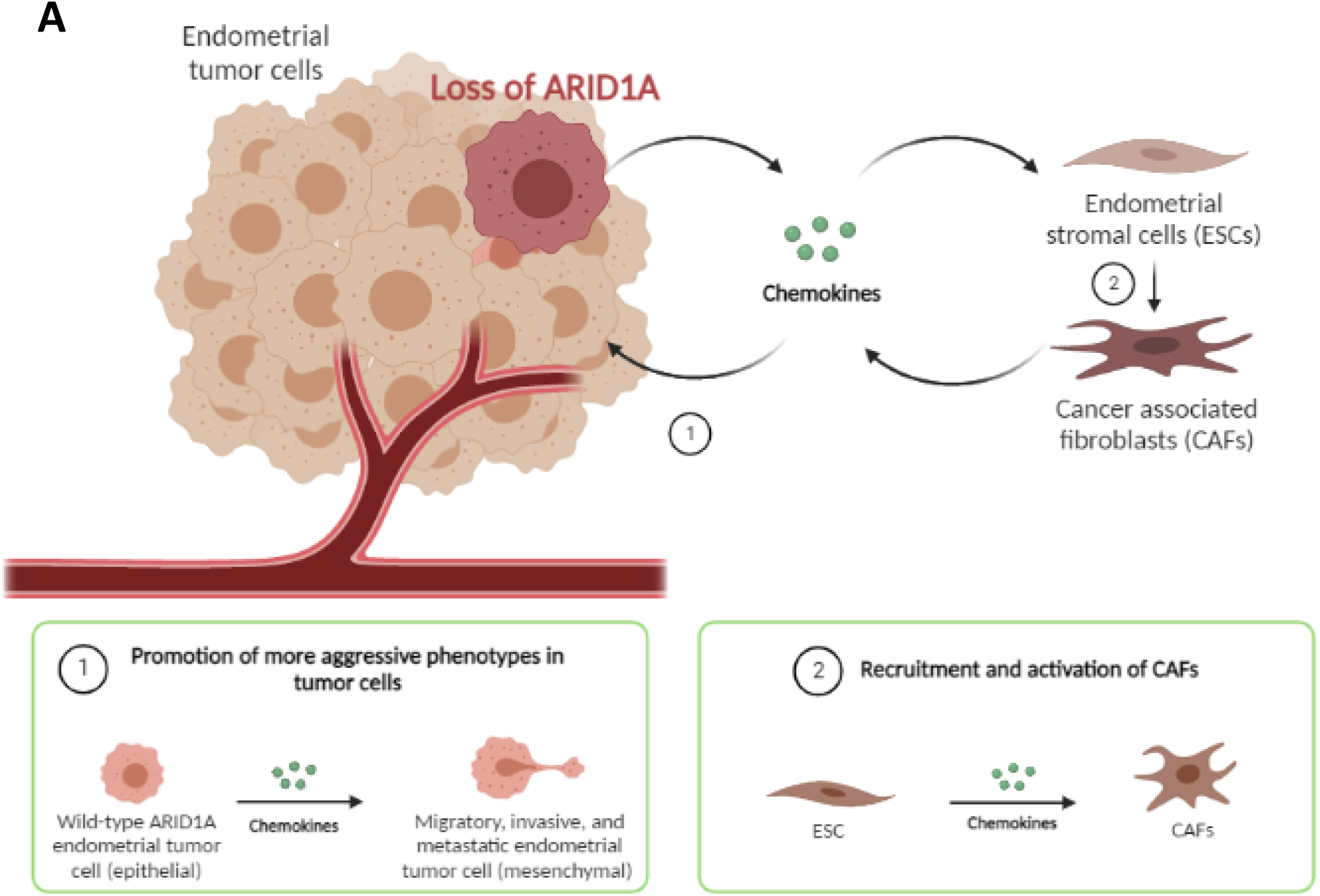
Schematic model proposed. Endometrial tumor cells with loss of ARID1A expression induce TME pro-tumoral reprograming by chemokine secretion. ARDI1A depleted endometrial tumour cells secreted high levels of chemokines. On one hand, these high levels of chemokines promote the acquisition of more aggressive behaviours in surrounding ARID1A wild-type endometrial tumor cells, inducing migration, invasion and dissemination. On the other hand, these enhanced levels of secreted chemokines promote the recruitment and activation of CAFs. Created with Biorender.com.

## Discussion

Tumors consist in a complex ecosystem composed not only of tumor cells, but also of stromal and immune cells, as well as soluble factors and components of the extracellular matrix^15^. Whitin this TME, several reciprocal interactions exist between the cells and with the rest of TME components, which play a fundamental role in the different parts of the tumor development and progression^26^. These interactions could be directly by cell-cell contact or by the secretion of molecules such as cytokines^26^. Local dissemination of the primary tumor through adjacent normal tissue is the initial step of metastasis, and tumor secreted molecules are crucial in this dissemination^26^. Several studies describe that within the primary tumor, tumor cells with more aggressive phenotypes are able to promote these phenotypes in surrounding primary tumor cells with less aggressive tumor phenotypes, promoting tumor progression and dissemination or the acquisition of resistance to antitumor therapies through the secretion of soluble factors^24,32^. Previously we showed that ARID1A loss in endometrial tumor cells promotes the activation of EMT programs and the acquisition of metastatic properties in these cells^23^. Here, we describe that ARID1A depleted endometrial tumor cells foster these acquired phenotypes in surrounding ARID1A wild-type endometrial tumor cells. Through *in vitro* assays employing CM and co-cultures models, we revealed the ability of endometrial tumor cells with loss of ARID1A to promote migration, invasion and EMT phenotypes in endometrial tumor cells with ARID1A normal expression, mainly through soluble factors secretion by local paracrine signals. Soluble factors secreted by tumor cells, in addition to playing an important role in tumor invasion at local level, also play an essential role in more advanced phases of the metastatic cascade such as intravasation, circulation, extravasation and metastatic colonization^14,26^. Our results demonstrated that signals from ARID1A depleted endometrial tumor cells are not only at local level, but that pro-tumoral signals are extrapolated at distant locations spread through the bloodstream, possibly playing a functional role in more advances stages of metastasis process that support the metastatic dissemination of ARID1A wildtype endometrial tumor cells. However, while the presented data indicate that soluble signals from ARID1A altered endometrial tumor are expanded from the primary TME, future studies will be necessary to stablish whether these signals promote pre-metastatic niches formation in secondary organs.

CAFs are the major component of the TME stroma and play a crucial role in tumor progression and aggressiveness^15,16^. Several works have demonstrated that tumor cells directly modulate activation and pro-tumoral function of CAFs through the secretion of paracrine signals^26,33–35^. Once activated, these CAFs promote tumor cells aggressiveness by secretion of soluble factors, generating a reciprocal interrelationship between tumor cells and CAFs Our data show that ARID1A depleted endometrial tumor cells induce ESC activation. We demonstrate by *in vitro* assays that these cells secrete signals that induce the activation and pro-tumoral activities of ESC, acquiring CAF phenotypes. Specifically in endometrium, where the stromal component is very abundant and is found evolving the epithelial glands, stromal cells play a very important role in the growth, communication and differentiation of epithelial cells^17,18^. Supported by our results, this suggest that in EC, ARID1A may play a key role in TME regulation and tumor progression. Given TME changes are more conserved between tumors, therapeutic strategies based on TME could be of great importance for the treatment of endometrial tumors with loss of ARID1A expression.

Finally, our data show that loss of ARID1A in endometrial tumor cells leads to alteration of the tumor secretory profile. Specifically, by analysis of CM from our *in vitro* models, we demonstrate a prominent significative increase in several chemokine levels, identifying high chemokine secretion as a pro-tumoral characteristic of EC cells with ARID1A alterations. Overall, our data suggest that in ECs with loss of ARID1A, increased chemokine secretion is responsible for generating a positive feedback loop that reprograms a reactive TME (Fig. 6). Indeed, supported by our *in vivo* data, we also propose that chemokine levels are increased in plasma from mice with ARID1A depleted endometrial tumors. The strong implication of chemokines in the metastatic dissemination process has been described in a wide range of solid tumors^26^. In particular, a study of Brile Chung et al. showed that in breast cancer, chemokines plays a critical role in the establishment of pre-metastatic niches facilitating tumor cell dissemination and metastatic formation in brain^36^. Taken together, this supports the hypothesis of ARID1A may play a key role in TME regulation and EC dissemination.

In sum, the data presented here demonstrate a potent novel role for ARID1A in EC stablishing a pro-tumorigenic environment and promoting metastatic dissemination by chemokine secretion in EC. We propose that chemokine signaling disruption strategies could be an exciting therapeutic possibility in high-grade/stage, invasive or metastatic EC harboring ARID1A mutations, since metastasis is the main cause of death in patients with cancer.

## Abbreviations

3D: three dimensions
ARID1A: AT-rich interactive domain-containing protein 1A
CAF: cancer associated fibroblast
CM: conditioned media
Cre:ERT: tamoxifen-inducible cre estrogen receptor recombinase
DNA: deoxyribonucleic acid
EC: endometrial cancer
ECM: extracellular matrix
EEC: endometrioid endometrial cancer
EGFP: enhanced green fluorescent protein
EMT: epithelial–mesenchymal transition
ESC: endometrial stromal cell
EV: empty vector
GFP: green fluorescent protein
IL: interleukin
IK: Ishikawa
MMP: matrix metalloproteinase
NEEC: non-endometrioid endometrial cancer
OCCC: ovarian clear cell carcinoma
PTEN: phosphatase and tensin homolog
RNA: ribonucleic acid
sgRNA: single guide RNA
shRNA: short hairpin RNA
SNAIL1: snail family transcriptional repressor 1
SPF: specific pathogen-free
SWI/SNF: switch/Sucrose non-fermentable
TGFb: transforming growth factor beta
TME: tumor microenvironment
VEGF: Vascular endothelial growth factor
WT: wild-type
ZEB: zinc finger E-box-binding homeobox 1

## Acknowledgements

This work has been funded by the Instituto de Salud Carlos III (ISCIII) through the projects PI20/00502, CB16/12/00231, (Cofounded by European Regional Development Fund (ERDF) “a way to make Europe” and ESF “Investing in your future). We also thank the Grups consolidats de la Generalitat de Catalunya (2017SGR1368) and the Asociación Española Contra el Cáncer (AECC; Grupos Estables 2018: GCTRA1804MATI). C. M-L. holds a predoctoral fellowship “Ajuts 2021 de Promoció de la Recerca en Salut-9^a^ edició” from IRBLleida/Diputació de Lleida. We also want to thank the CERCA programme/Generalitat de Catalunya for institutional support.

## Data Availability

The data that support the findings of this study are available on request from the corresponding author.

## Authors’ contributions

Study conception and design: M-L. C.; E.N.; M-G. X. Acquisition and interpretation of data: M-L. C.; E.N. All authors were involved in the writing of the manuscript and gave the submitted version their final approval.

## Supporting Information

**Figure supplementary 1:**
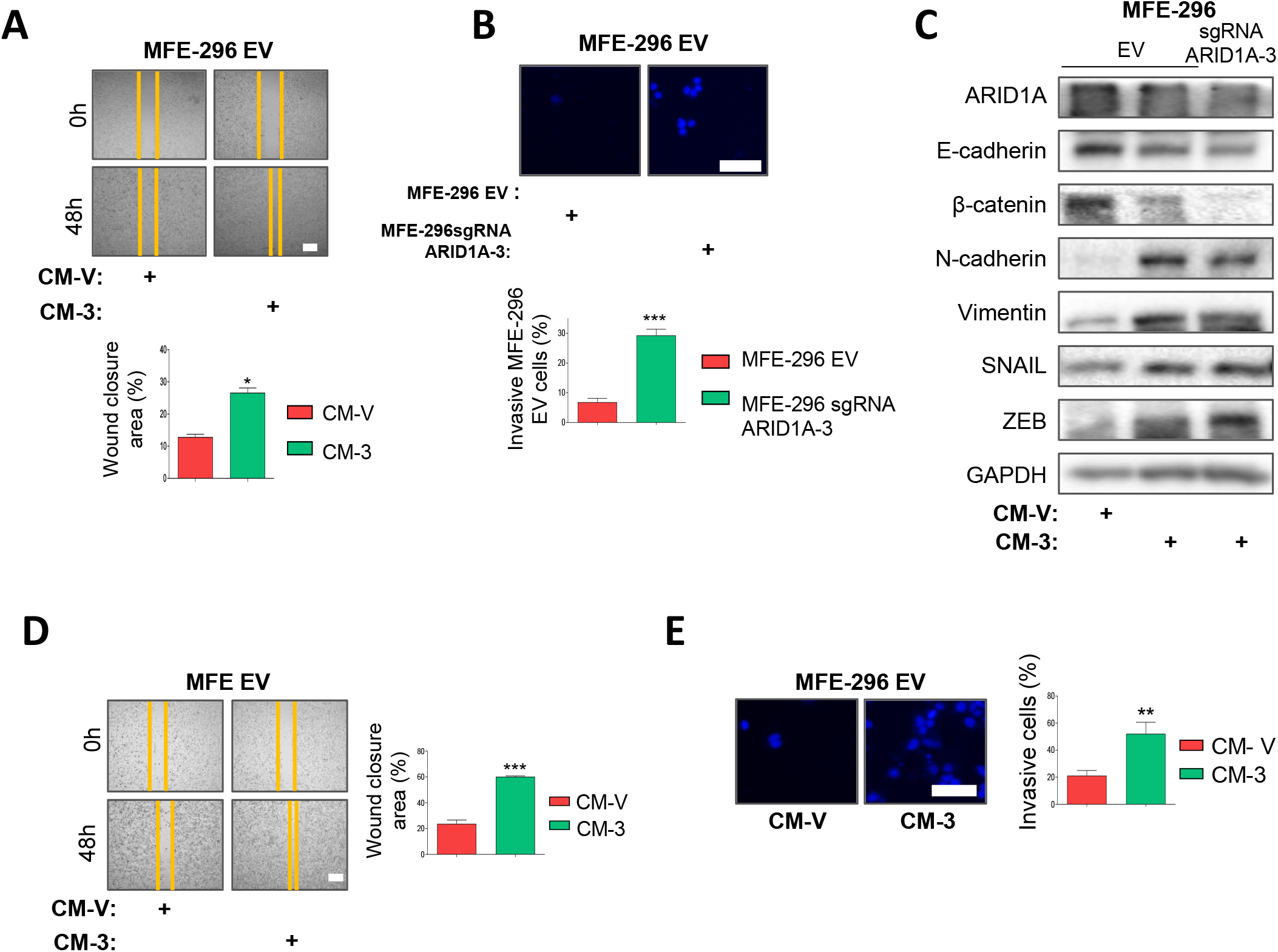
**A)** Representative images at time 0 and 48 h after scratch, of wound-healing assay performed in IK cells infected with the EV treated with conditioned media collected from MFE-296 cells infected with lentiviruses carrying sgRNA against *ARID1A* (lentiCRISPRv2-ARID1A-3) (CM-3) or EV (CM-V). Bottom blot shows quantification of wound closure area between the indicated time. Scale bars: 200 μm. **B)** Representative images of nuclear Hoechst staining of transwell invasion assay after the cotton swab in MFE-296 control cells chemoattract by MFE-296 cells infected with lentiviruses carrying sgRNA against *ARID1A* (lentiCRISPRv2-ARID1A-3) or EV for 48 hours (upper panel), and quantification of Matrigel® invasive cells (bottom plot). Scale bars: 50 μm. **C)** Western blot analysis of ARID1A, E-cadherina, β-catenina, N-cadherina, vimentina, SNAIL and ZEB in MFE-296 cells infected with the EV treated with CM-3 or CM-V for 48 hours. GAPDH was used as a loading control. **D)** Representative images at time 0 and 48 h after scratch, of wound-healing assay performed in MFE-296 cells infected with the EV previously treated for 48h with conditioned media collected from IK or MFE-296 cells infected with lentiviruses carrying sgRNA against *ARID1A* (lentiCRISPRv2-ARID1A-3) (CM-3) or EV (CM-V). Right blots show quantification of wound closure area between the indicated time. Scale bars: 200 μm. **E)** Representative images of nuclear Hoechst staining of transwell invasion assay after the cotton swab in IK EV cells pretreated for 48 with conditioned media collected from MFE-296 cells infected with lentiviruses carrying sgRNA against *ARID1A* (lentiCRISPRv2-ARID1A-3) (CM-3) or EV (CM-V) (left panel), and quantification of Matrigel® invasive cells (right plot). Scale bars: 50 μm. Graph values are the mean and error bars represented as mean ± S.E.M. Statistical analysis was performed using unpaired 2-tailed *Student t-test* analysis. * *p* < 0.05; \*\*\**p* < 0.001.

**Figure supplementary 2:**
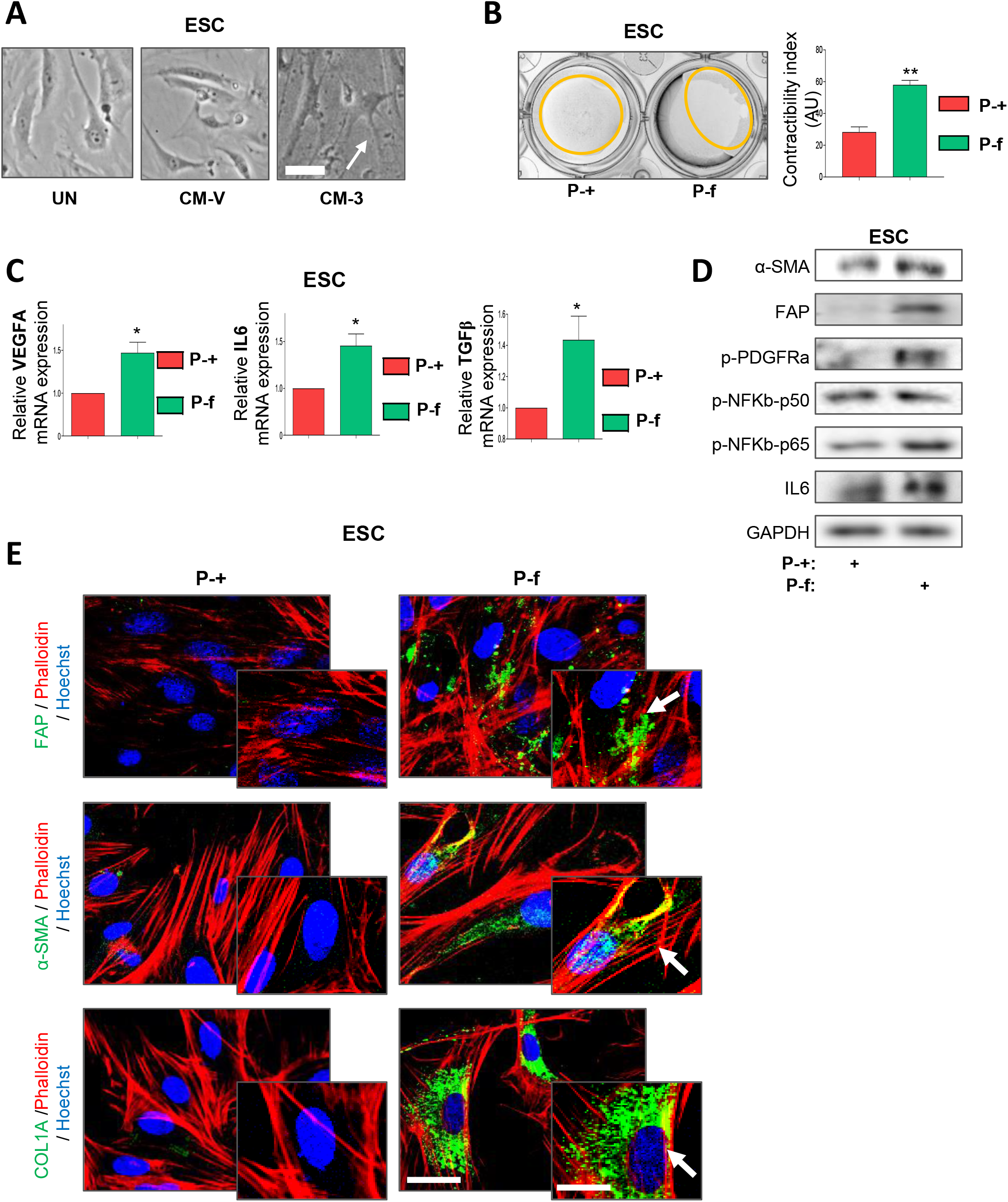
**A)** Representative phase contrast images of live cell morphologies of ESC treated for 48h with CM collected from IK cells infected with lentivirus carrying sgRNA against ARID1A (lentiCRISPRv2 ARID1A-3) (CM-3) or empty vector (CM-V). **B)** Upper, representative macroscopic images of collagen gels remodelled by ESC previously treated for 48h with plasma collected from *Cre:ERT; Pten f/f; Arid1a +/+* (P-+) or *Cre:ERT; Pten f/f; Arid1a f/f* (P-f) mice with endometrial tumors. Bottom, histogram representing the quantification of the contractility index **C)** RT-qPCR of *Vegfa, Il6* and *Tgfβ1* mRNA expression of ESC treated with P-f or P-+ for 48h. **D)** Representative immunoblotting images of α-SMA, FAP, p-PDGFRα (Tyr754), p-NFκβ-p50 (Ser337), p-NFκβ-p65 (Ser536) and IL6 protein expression in ESC treated with P-f or P-+ for 48h. GAPDH was used as a loading control. **E)** Representative immunofluorescence images of the stromal activation markers FAP, α-SMA and COL1A in ESC treated with P-f or P-+ for 48h. Phalloidin and Hoechst were used as actin cytoskeleton and nucleus markers respectively. Graph values are the mean and error bars represented as mean ± S.E.M. Statistical analysis was performed using unpaired 2-tailed *Student t-test* analysis. * *p* < 0.05; \*\*\**p* < 0.001.

## References

1. Sung H, Ferlay J, Siegel RL, et al. Global Cancer Statistics 2020: GLOBOCAN Estimates of Incidence and Mortality Worldwide for 36 Cancers in 185 Countries. CA Cancer J Clin. 2021;71(3):209–249. doi:10.3322/CAAC.21660

2. Constantine GD, Kessler G, Graham S, Goldstein SR. Increased incidence of endometrial cancer following the women’s health initiative: An assessment of risk factors. J Women’s Heal. 2019;28(2):237–243. doi:10.1089/JWH.2018.6956/ASSET/IMAGES/LARGE/FIGURE3.JPEG

3. Bokhman J V. Two pathogenetic types of endometrial carcinoma. Gynecol Oncol. 1983;15(1):10–17. doi:10.1016/0090-8258(83)90111-7

4. Matias-Guiu X, Prat J. Molecular pathology of endometrial carcinoma. Histopathology. 2013;62(1):111–123. doi:10.1111/his.12053

5. Suarez AA, Felix AS, Cohn DE. Bokhman Redux: Endometrial cancer “types” in the 21st century. Gynecol Oncol. 2017;144(2):243–249. doi:10.1016/J.YGYNO.2016.12.010

6. Piulats JM, Guerra E, Gil-Martín M, et al. Molecular approaches for classifying endometrial carcinoma. Gynecol Oncol. 2017;145(1):200–207. doi:10.1016/J.YGYNO.2016.12.015

7. Getz G, Gabriel SB, Cibulskis K, et al. Integrated genomic characterization of endometrial carcinoma. Nature. 2013;497(7447):67–73. doi:10.1038/nature12113

8. Mullen J, Kato S, Sicklick JK, Kurzrock R. Targeting ARID1A mutations in cancer. Cancer Treat Rev. 2021;100:102287. doi:10.1016/j.ctrv.2021.102287

9. Megino-Luque C, Sis P, Mota-Martorell N, et al. ARID1A-deficient cells require HDAC6 for progression of endometrial carcinoma. Mol Oncol. 2022;16:2235. doi:10.1002/1878-0261.13193

10. Xu S, Tang C. The Role of ARID1A in Tumors: Tumor Initiation or Tumor Suppression? Front Oncol. 2021;11:3891. doi:10.3389/FONC.2021.745187/BIBTEX

11. Mao TL, Shih IM. The roles of ARID1A in gynecologic cancer. J Gynecol Oncol. 2013;24(4):376. doi:10.3802/JGO.2013.24.4.376

12. Suryo Rahmanto Y, Shen W, Shi X, et al. Inactivation of Arid1a in the endometrium is associated with endometrioid tumorigenesis through transcriptional reprogramming. Nat Commun. 2020;11(1). doi:10.1038/S41467-020-16416-0

13. Gibson WJ, Hoivik EA, Halle MK, et al. The genomic landscape and evolution of endometrial carcinoma progression and abdominopelvic metastasis. Nat Genet. 2016;48(8):848–855. doi:10.1038/ng.3602

14. Bejarano L, Jordāo MJC, Joyce JA. Therapeutic Targeting of the Tumor Microenvironment. Cancer Discov. 2021;11(4):933–959. doi:10.1158/2159-8290.CD-20-1808

15. Anderson NM, Simon MC. The tumor microenvironment. Curr Biol. 2020;30(16):R921–R925. doi:10.1016/J.CUB.2020.06.081

16. Sahai E, Astsaturov I, Cukierman E, et al. A framework for advancing our understanding of cancer-associated fibroblasts. Nat Rev Cancer. 2020;20(3):174–186. doi:10.1038/s41568-019-0238-1

17. Sahoo SS, Zhang XD, Hondermarck H, Tanwar PS. The Emerging Role of the Microenvironment in Endometrial Cancer. Cancers (Basel). 2018;10(11). doi:10.3390/CANCERS10110408

18. Casas-Arozamena C, Abal M. Endometrial Tumour Microenvironment. Adv Exp Med Biol. 2020;1296:215–225. doi:10.1007/978-3-030-59038-3_13

19. Eritja N, Llobet D, Domingo M, et al. A novel three-dimensional culture system of polarized epithelial cells to study endometrial carcinogenesis. Am J Pathol. 2010;176(6):2722–2731. doi:10.2353/AJPATH.2010.090974

20. Ranftl RE, Calvo F. Analysis of breast cancer cell invasion using an organotypic culture system. In: Methods in Molecular Biology. Vol 1612. Humana Press Inc.; 2017:199–212. doi:10.1007/978-1-4939-7021-6_15

21. Gao X, Tate P, Hu P, Tjian R, Skarnes WC, Wang Z. ES cell pluripotency and germ-layer formation require the SWI/SNF chromatin remodeling component BAF250a. Published online 2008. Accessed August 25, 2022. https://www.pnas.org/cgi/content/full/

22. Mirantes C, Eritja N, Dosil MA, et al. An inducible knockout mouse to model the cell-autonomous role of PTEN in initiating endometrial, prostate and thyroid neoplasias. DMM Dis Model Mech. 2013;6(3):710–720. doi:10.1242/DMM.011445/258755/AM/AN-INDUCIBLE-KNOCK-OUT-MOUSE-TO-MODEL-CELL

23. Megino-Luque C, Sisó P, Mota-Martorell N, et al. ARID1A-deficient cells require HDAC6 for progression of endometrial carcinoma. Mol Oncol. Published online March 2, 2022. doi:10.1002/1878-0261.13193

24. Obenauf AC, Zou Y, Ji AL, et al. Therapy-induced tumour secretomes promote resistance and tumour progression. Nature. 2015;520(7547):368–372. doi:10.1038/NATURE14336

25. Paltridge JL, Belle L, Khew-Goodall Y. The secretome in cancer progression. Biochim Biophys Acta. 2013;1834(11):2233–2241. doi:10.1016/J.BBAPAP.2013.03.014

26. Hussain S, Peng B, Cherian M, Song JW, Ahirwar DK, Ganju RK. The Roles of Stroma-Derived Chemokine in Different Stages of Cancer Metastases. Front Immunol. 2020;11:1. doi:10.3389/FIMMU.2020.598532

27. Pereira BA, Vennin C, Papanicolaou M, et al. CAF Subpopulations: A New Reservoir of Stromal Targets in Pancreatic Cancer. Trends in Cancer. 2019;5(11):724–741. doi:10.1016/J.TRECAN.2019.09.010

28. Sahai E, Astsaturov I, Cukierman E, et al. A framework for advancing our understanding of cancer-associated fibroblasts. Nat Rev Cancer 2020 203. 2020;20(3):174–186. doi:10.1038/s41568-019-0238-1

29. Lee BY, Hogg EKJ, Below CR, et al. Heterocellular OSM-OSMR signalling reprograms fibroblasts to promote pancreatic cancer growth and metastasis. Nat Commun 2021 121. 2021;12(1):1–20. doi:10.1038/s41467-021-27607-8

30. Avgustinova A, Iravani M, Robertson D, et al. Tumour cell-derived Wnt7a recruits and activates fibroblasts to promote tumour aggressiveness. Nat Commun 2016 71. 2016;7(1):1–14. doi:10.1038/ncomms10305

31. Ferrari N, Ranftl R, Chicherova I, et al. Dickkopf-3 links HSF1 and YAP/TAZ signalling to control aggressive behaviours in cancer-associated fibroblasts. Nat Commun. 2019;10(1). doi:10.1038/s41467-018-07987-0

32. Celià-Terrassa T, Bastian C, Liu D, et al. Hysteresis control of epithelial-mesenchymal transition dynamics conveys a distinct program with enhanced metastatic ability. Nat Commun 2018 91. 2018;9(1):1–12. doi:10.1038/s41467-018-07538-7

33. Araujo AM, Abaurrea A, Azcoaga P, et al. Stromal oncostatin M cytokine promotes breast cancer progression by reprogramming the tumor microenvironment. J Clin Invest. 2022;132(7). doi:10.1172/JCI148667

34. Avgustinova A, Iravani M, Robertson D, et al. Tumour cell-derived Wnt7a recruits and activates fibroblasts to promote tumour aggressiveness. Nat Commun 2016 71. 2016;7(1):1–14. doi:10.1038/ncomms10305

35. Liu J, Chen S, Wang W, et al. Cancer-associated fibroblasts promote hepatocellular carcinoma metastasis through chemokine-activated hedgehog and TGF-β pathways. Cancer Lett. 2016;379(1):49–59. doi:10.1016/J.CANLET.2016.05.022

36. Chung B, Esmaeili AA, Gopalakrishna-Pillai S, et al. Human brain metastatic stroma attracts breast cancer cells via chemokines CXCL16 and CXCL12. npj Breast Cancer 2017 31. 2017;3(1):1–9. doi:10.1038/s41523-017-0008-8

